# Flexible Learning-Free Segmentation and Reconstruction for Sparse Neuronal Circuit Tracing

**DOI:** 10.1101/278515

**Authors:** Ali Shahbazi, Jeffery Kinnison, Rafael Vescovi, Ming Du, Robert Hill, Maximilian Joesch, Marc Takeno, Hongkui Zeng, Nuno Macarico da Costa, Jaime Grutzendler, Narayanan Kasthuri, Walter J. Scheirer

## Abstract

Imaging is a dominant strategy for data collection in neuroscience, yielding stacks of images that often scale to gigabytes of data for a single experiment. Machine learning algorithms from computer vision can serve as a pair of virtual eyes that tirelessly processes these images, automatically constructing more complete and realistic circuits. In practice, such algorithms are often too error-prone and computationally expensive to be immediately useful. We address these shortcomings with a new fast, flexible, learning-free method for sparse segmentation and reconstruction of neural volumes. Unlike learning methods, our Flexible Learning-free Reconstruction of Imaged Neural volumes (FLoRIN) pipeline exploits structure-specific contextual clues and requires no training. This approach generalizes across different modalities, including serially-sectioned scanning electron microscopy (sSEM) of genetically labeled and contrast enhanced processes, spectral confocal reflectance (SCoRe) microscopy, and high-energy synchrotron X-ray microtomography (μCT) of large tissue volumes. We deploy the FLoRIN pipeline on newly published and novel mouse datasets, demonstrating the high biological fidelity of the pipeline’s reconstructions, which are of sufficient quality for preliminary biological study. Compared to existing supervised learning methods, it is both significantly faster (up to several orders of magnitude) and produces high-quality reconstructions that are robust to noise and artifacts.

## Introduction

The ability to comprehensively study the brain is within reach. Advances in tissue preparation and imaging technologies have allowed researchers to collect vast amounts of targeted structural data, potentially enabling new multi-resolution, multi-modal studies of neural volumes. Brain imaging techniques from microscopy such as serially-sectioned scanning electron microscopy (sSEM), high-energy synchrotron X-ray microtomography (μCT), and spectral confocal reflectance (SCoRe) microscopy provide high-quality images with nanoscale or single-neuron resolution. To study these volumes, however, it is necessary to find the regions of interest (segmentation) and properly identify segmented structures (reconstruction).

There are two strategies to reconstruct a microscopic volume of the brain: segmenting only specific microstructures (sparse segmentation) or segmenting all structures contained within this volume (dense segmentation). Ultimately, we would like to apply dense segmentation to all neural volumes, however it is a time-consuming, error-prone process^1^. Additionally, in many cases it is not necessary to densely segment and reconstruct a neural volume to gain valuable insight: μ CT X-ray volumes contain a relatively small portion of microstuctures, however they can offer insight into the properties and distributions of cells in different regions of the brain. SCoRe imaging, too, often focuses on single cells and axons at a time that must be distinguished from background noise. In these cases, a sparse segmentation scheme is sufficient to begin studying the imaged region and requires less fine-tuning than its dense counterpart. Solving the sparse segmentation and reconstruction problem is the focus of this study.

To segment and analyze this variety of microscopy techniques, different methods have been employed from purely manual segmentation to complicated multi-layer deep convolutional neural networks. Each method offers different types of dense and sparse segmentation at a different time/quality cost. Despite all the differences between manual annotation methods and learning-based methods in speed, accuracy and efficiency, both of them need expert labor to annotate stacks of ground truth. This slows down the process of studying a new sample, new imaging output, or training a basic machine learning or deep learning approach.

With respect to approaches for generating manual annotations, a number of software frameworks are available, including Eyewire^2^, D2P^3^, Mojo^4^, Catmaid^5^, KNOSSOS^6^, Ssecrett^7^, TrakEM2^8^ and ITK-SNAP^9^. Some neuronal circuit tracing studies are based on exhaustive manual proof-reading of results from an algorithm^10,11^, which, like the creation of ground-truth data, is not practical for datasets consisting of high resolution images or large volumes. For example, Takemura *et al*.^10^ described how the reconstruction of a relatively small volume of *Drosophila* brain required ~ 12,940 person-hours of proof-reading.

To overcome the limitations of manual annotation, expert-verified reconstructions may be used to train machine learning models to perform automatic segmentation and reconstruction. A wide variety of methods — highly dependent on supervised machine learning — exist for the dense EM segmentation and reconstruction problem. The methods include SVM-based algorithms^12-16^, Random Forests^12,17-24^, Conditional Random Fields^22^, and Artificial Neural Networks^25-30^ (i.e., deep learning). These machine learning approaches can be found in popular software packages for connectomics image analysis such as Rhoana^31^ and Ilastik^18,19,32^. In addition to traditional artificial neural network-based approaches, research into fully-convolutional networks for image segmentation related to microscopy produced the U-Net^33^ architecture. This type of network uses convolutions, downsampling, and upsampling in a U-shaped configuration to home in on features and good candidate output segmentations. Additionally, intermediate results from the downsampling side of the network are fed to the corresponding convolutional units on the upsampling side to provide extra information to later layers. These networks have been applied to both the dense^33,34^ and sparse^34,35^ segmentation problems in the 2D and 3D domains.

In spite of these machine learning innovations, neuroscientists must remain in-the-loop when applying deep learning methods to provide high-quality training data. Images from neuroscience experiments are complex, containing a wide variety of cells and other micro-structures that may only be reliably and correctly identified by expert annotators. This is not a good scenario for deep learning methods, which require a prohibitively large amount of annotated training data.

Even if an image volume has been annotated, deep learning methods require a vast amount of specialized computational resources. Artificial neural networks can take days or weeks to train, and feature extraction, while faster than training, is still inefficient even on a graphical processing unit (GPU). For just 100 images with a resolution of 1024 × 1024, a state-of-the-art network for connectomics took 59 hours for training and evaluation^29^. Dong *et al*.^36^ described spending three days training a CNN to sparsely segment zebrafish cells in a stack of 100 1024 × 1344 images. Ciresan et al.^28^ trained a 4-layer artificial neural network on the 2012 ISBI challenge dataset over the course of several days, with each training epoch taking between 170-340 minutes. Similarly, Teikari *et al.*^37^ reported training their VD2D3D convolutional net to segment vasculature in multiphoton microscopy for 24 days. Such timescales are unavoidable as artificial neural networks become deeper and more complex and as datasets increase in size.

In addition to long training times, learning-based methods are prone to dataset bias, limiting their generality. This bias can be caused by a low training set to testing set size ratio^38^, overfitting^39^, or insufficient training data^40^. The latter problem is not uncommon in neuroscience, where expert manual annotations tend to be expensive to create.

Given the current limitations of deep learning for neural circuit reconstruction, and until more expert annotations are available to better train learning methods, it is worth re-evaluating the current assumption that machine learning should be indiscriminately applied to all pattern recognition problems. New tissue preparations and imaging techniques have drastically improved the signal-to-noise ratio across imaging modalities, thus automated segmentation and reconstruction methods for neural circuit tracing can now look to reduce manual and computational overhead by exploiting geometric and grayscale intensity cues with classical image processing, removing training entirely.

Here we introduce the Flexible Learning-free Reconstruction of Imaged Neural volumes pipeline (FLoRIN) for automatic segmentation and reconstruction (see: Methods). In contrast with supervised and unsupervised machine learning methods, FLoRIN segments and reconstructs neural volumes using classical, learning-free computer vision techniques in 2D or 3D. Figure 1 provides an overview of the FLoRIN pipeline. This method distinguishes itself from other segmentation and reconstruction efforts by a number of facets:

- **A 3-Stage Pipeline**. FLoRIN starts with raw images and processes them in three stages: Segmentation, which binarizes regions of interest; Identification, which removes artifacts and labels microstrutures; and Reconstruction, which outputs 3D volumes and statistical data. During each stage, a flexible set of learning-free computer vision techniques may be applied to adapt to the challenges of datasets across many modalities.
- **A Novel Thresholding Technique**. During the Segmentation stage, we apply our novel thresholding algorithm, N-Dimensional Neighborhood Thresholding (NDNT), to quickly and accurately find microstructures in images. NDNT is an extension of the method introduced by Bradley and Roth^41^ into N-dimensions based on the formulas presented by Tapia^42^.
- **Mechanisms for Studying Microstructures**. In addition to labeled images, FLoRIN creates a database with the geometric and grayscale properties of detected microstructures. This database may be used to compute statistics about the reconstructed volume, including cell count, distribution of cell sizes, vasculature length, and more.

**Figure 1.**
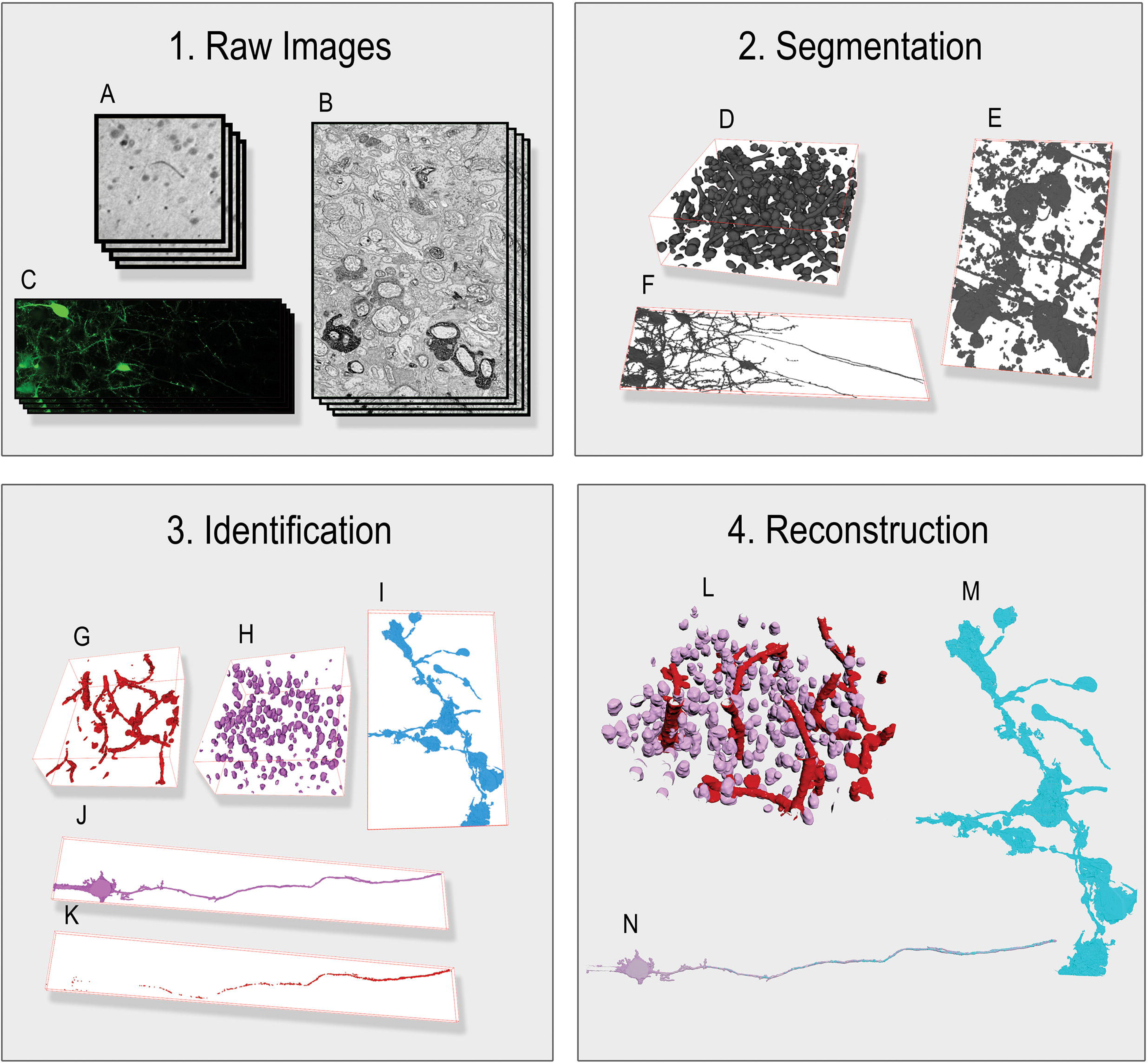
The Flexible Learning-free Reconstruction of Imaged Neural volumes (FLoRIN) pipeline is software for sparse segmentation and reconstruction of neural volumes. Unlike current methods that rely on learned models, FLoRIN uses classical learning-free image processing and computer vision methods to quickly and effectively segment and reconstruct microstructures in microscopic images. The FLoRIN pipeline consists of the following steps: (1) Raw images are loaded into the pipeline. (2) Images are segmented using N-Dimensional Neighborhood Thresholding to binarize structures of interest. (3) Connected components in the binarized images are filtered to remove noise and distinguish between classes of microstructures. (4) Each class of microstructure is saved in its own volume and a statistical report is generated based on the location, geometry, and grayscale histogram of each individual structure. By virtue of being learning-free, FLoRIN is generalizable across multiple imaging modalities, including μ CT X-ray (A, D, G, H, L), sSEM (B, E, I, M), and SCoRe (C, F, J, K, N) images.

FLoRIN is dramatically faster than deep learning methods, more tolerant to noise and image artifacts, and generalizable across datasets and modalities. The novel NDNT algorithm also outperforms other thresholding methods at the sparse segmentation task. To demonstrate the improvements made possible by FLoRIN, we segment and reconstruct two μ CT X-ray volumes, two sSEM volumes, and one SCoRe volume. Where ground-truth annotations are available, we compare the results of FLoRIN against 3D U-Net^34^, a state-of-the-art fully convolutional neural network designed to segment neural volumes. In every case, FLoRIN produces higher-quality reconstructions than 3D U-Net in a fraction of the time without training. NDNT segmentations are also shown to be of significantly higher quality than those of standard thresholding methods. We conclude that our pipeline is suitable for sparse segmentation and reconstruction of both *in-vivo* and *ex-vivo* imaging modalities.

## Results

We applied FLoRIN to sparsely segment and reconstruct neural volumes from three different modalities: μ CT X-ray, sSEM, and SCoRe. Additionally, we evaluated each volume with a number of standard thresholding methods (Supplementary Methods: Non-Machine Learning Segmentation Method Evaluation). Two of the image volumes were expert-annotated, and in these cases we also trained U-Net and 3D U-Net models (Supplementary Methods: U-Net & 3D U-Net Training) to compare FLoRIN against a state-of-the-art deep learning model designed for microscopic image segmentation.

### μ CT X-ray Volumes

X-ray computed tomography is an imaging technique that captures volumetric views of tissue samples. Since the contrast mechanism is directly related to the sample density, it is possible to distinguish between macrostructures like cells, myelinated bundles, and vasculature. We examine two such μ CT X-ray volumes imaged at micron-scale resolutions. In this study, two different μ CT image volumes were reconstructed: a standard rodent brain volume (SRB) with human-generated annotations and an APEX2-labeled rodent brain volume (ALRB) that includes APEX2-labeled cells.

### Standard Rodent Brain (SRB)

The SRB volume is relatively small, spanning only a 300 × 300 × 100 voxel region imaged at a resolution of 1.2μm^43^ (5.1 MB). But it is accompanied by a partial human annotation of the imaged cells, making it suitable as a benchmark for automatic segmentation and reconstruction methods. Cells and vasculature may be visually differentiated (Figure 2 Panels A & B), however automatic segmentation is complicated by grayscale shifts and cells that are within close proximity of each other, which can both lead to merge errors and voxel mis-classifications. Without careful consideration of these factors or extensive training, automated methods will fail to properly segment the cells and vasculature. With this in mind, we created reconstructions of the cells and vasculature in SRB using the full FLoRIN pipeline and a number of learning-based and learning-free segmentation methods.

**Figure 2.**
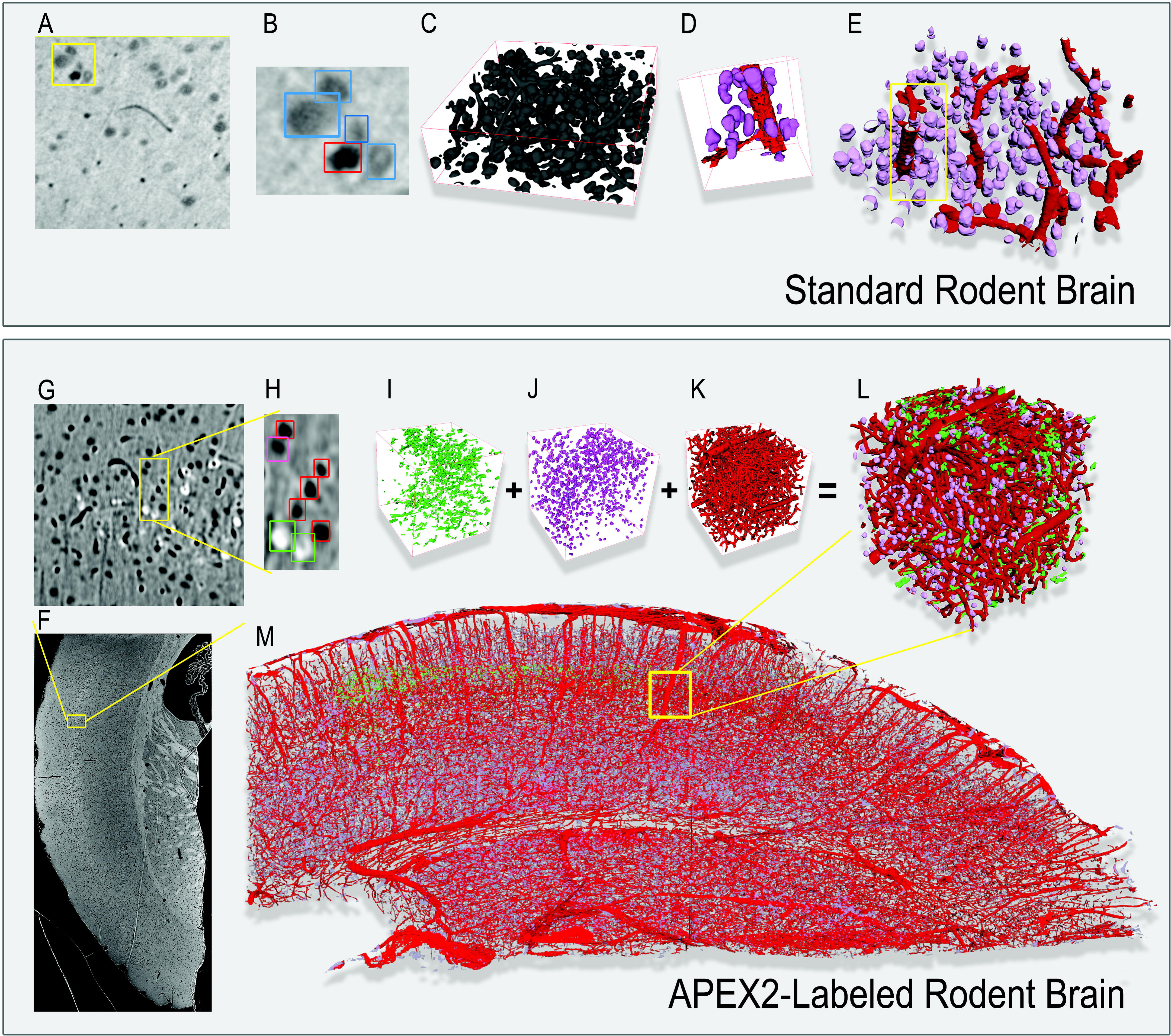
FLoRIN reconstructions of the Standard Rodent Brain (SRB) (top) and APEX2-labeled Rodent Brain sample (ALRB) (bottom) μ CT X-ray volumes. (A) Within the SRB volume, cells and vasculature are visually distinct in the raw images, with vasculature appearing darker than cells. (B) Individual structures may be extremely close (such as the cells and vasculature in this example), making reconstruction efforts prone to merge errors. (C) Segmentation by FLoRIN extracts the structures from the images with minor noise and artifacts that are much smaller than the structures of interest. (D) FLoRIN is able to separate nearby structures based on geometric and grayscale features. (E) After filtering, the final reconstruction distinguishes between cells (purple) and vasculature (red). (F) The ALRB volume spans a large portion of the rodent brain. (G, H) In addition to non-APEX cells and vasculature, ALRB contains APEX2-labeled cells that appear lighter than the background. FLoRIN segmented and filtered the APEX2-labeled cells (I), non-APEX cells (J), and vasculature (K), then reconstructed them into a single volume (L). (M) Throughout the ALRB volume, the vasculature was segmented as a single connected component, and FLoRIN discovered 123,424 non-APEX2-labeled cells and 1,524 APEX2-labeled cells.

Overall, FLoRIN was able to most closely reconstruct the cells in the volume, discovering all 313 manually-annotated cells. An additional seven cells were also discovered and verified, for a total of 320. As shown in Table 1, FLoRIN generated a reconstruction with a mean Hausdorff distance of 9.61 × 10^−1^, at least an order of magnitude closer to the ground-truth than any other deep learning or learning-free method evaluated. We also evaluated the raw segmentation results of NDNT operating in 2D and 3D, and both of these methods created segmentations closer to the ground-truth than any other method evaluated. In this case, we observed that NDNT was the best segmentation method, and the full FLoRIN pipeline was able to refine these results further. Of the other evaluated methods, only the Sauvola thresholding^44^, Triangle thresholding^45^, 2D U-Net^33^, and MGAC^46^ methods were able to approach the performance of NDNT.

**Table 1.** Performance comparison between FLoRIN, NDNT, U-Net models, and standard thresholding methods on the SRB and ALRB X-Ray volumes. The SRB volume contains 320 total cells. U-Net results are the mean performance of 20 randomly-initialized nets. Asterisks indicate methods applied to small 3D blocks of the volume at a time. U-Net cell counts are reduced due to evaluation on the test set, a subset of 10 images.

| Methods | SRB |  |  | ALRB |
| --- | --- | --- | --- | --- |
|  | Mean Hausdorff Distance | Runtime (Minutes) | Cell Count | Runtime (Minutes) |
| **FLoRIN** | 0.9610 | 14.9111 | 320 | 45.17 |
| **NDNT\*** | 3.9076 | 7.8980 | 253 | 41.15 |
| **NDNT** | 5.6836 | 0.2981 | 220 | 2.77 |
| **Isodata\***<sup>49</sup> | 56.1870 | 0.9129 | 9 | 3.85 |
| **Isodata**<sup>49</sup> | 20.1399 | 0.0523 | 131 | 3.85 |
| **Li\***<sup>50</sup> | 83.5557 | 6.3752 | 4 | 35.84 |
| **Li**<sup>50</sup> | 18.4453 | 0.6358 | 140 | 1.97 |
| **Local**<sup>54</sup> | 58.4740 | 0.4692 | 7 | 7 |
| **MGAC**<sup>46</sup> | 10.5529 | 237.7531 | 111 | 1834.63 |
| **Niblack**<sup>53</sup> | 43.9941 | 1.3723 | 24 | 6.17 |
| **Otsu\***<sup>47</sup> | 100.5323 | 1.0140 | 3 | 4.47 |
| **Otsu**<sup>47</sup> | 20.5289 | 0.0576 | 126 | 0.48 |
| **Otsu 3D**<sup>47</sup> | 91.7326 | 0.0232 | 0 | 0.21 |
| **Sauvola**<sup>44</sup> | 6.2682 | 1.3383 | 198 | 6.22 |
| **Triangle\***<sup>45</sup> | 23.8607 | 0.8932 | 115 | 4.44 |
| **Triangle**<sup>60</sup> | 8.2826 | 0.0453 | 203 | 0.40 |
| **Yen\***<sup>52</sup> | 27.7905 | 0.8809 | 64 | 3.77 |
| **Yen**<sup>52</sup> | 9.6325 | 0.0460 | 175 | 0.46 |
| **U-Net**<sup>33</sup> | 9.618 ± 1.02<br>(for 10 slices) | Train: 145.891<br>Test: 0.049 | 65<br>(out of 79) | Not Applicable |
| **3D U-Net**<sup>34</sup> | 14.828 ± 2.66<br>(for 10 slices) | 0.0460 | 53<br>(out of 79) | Not Applicable |

As shown in Figure 2, the FLoRIN reconstruction captures the morphology of the cells and vasculature well. In Panels D & E of Figure 2, the cells are all smooth, localized structures while the vasculature segments weave and branch through the volume as connected tube-like structures. When superimposed in 2D (Supplementary Figure S3), the segmented areas visually correspond well with the original images, contrasting with the manual annotations which only capture a subset of cells and are (surprisingly) not faithful to the visible structures in the images. Despite the fact that the expert annotations only capture cells in SRB, both FLoRIN methods segmented both cells and vasculature simultaneously. U-Net^33^ and 3D U-Net^34^ could not be trained on vasculature in this volume and only segmented the cells. FLoRIN is thus able to extract useful information from images that human annotators may exclude or overlook due to time constraints.

FLoRIN was not the fastest method. While FLoRIN was able to segment and reconstruct SRB in 14.9 minutes, it was slower than all other thresholding methods. As noted, however, the reconstruction was much closer to the ground-truth than any other method, meaning that FLoRIN trades a modest amount of running time for significantly better results. Notably, FLoRIN was able to generate a reconstruction an order of magnitude faster than U-Net and 3D U-Net were able to train on SRB. While the deep learning methods can make predictions as quickly as the learning-free thresholding methods, doing so requires a trained model which may not be readily available.

Qualitative results of various methods are shown in Figure 3. As can be seen, the FLoRIN result includes full cells and only small amounts of noise. No vasculature is included in the reconstruction, which is as intended when attempting to match the manual annotations. Other methods either severely over-(as in 3D U-Net^34^, Local, and Otsu^47^) or under-segment (as in MGAC^46^) the image. The next best result after FLoRIN’s reconstruction and the NDNT segmentations, Sauvola^44^, includes artifacts and vasculature that NDNT was more robust to, which ultimately increased its Hausdorff distance to the ground-truth. As will be seen in the sSEM results, however, Sauvola’s method does not generalize to other modalities, while NDNT is able to do so.

**Figure 3.**
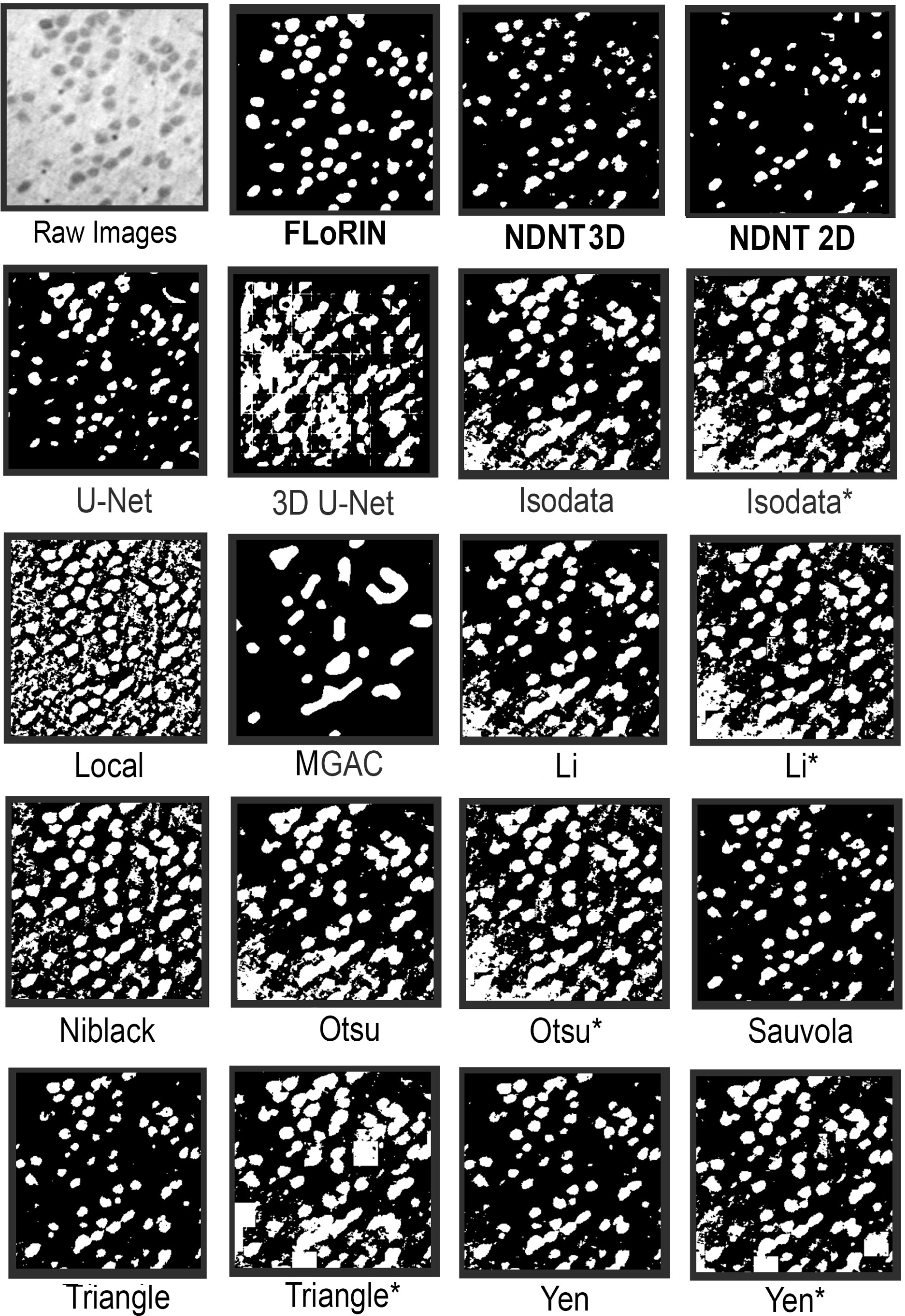
An example SRB image and the corresponding cell segmentation results of various methods. The FLoRIN reconstruction based on 3D NDNT, along with the raw 3D NDNT segmentation, captures all of the cells with only minor merge errors. All other methods severely over- or under-segment the image, and are susceptible to the grayscale shift in the lower-left corner of the image. The only other methods to come close to the FLoRIN results are Sauvola^44^ and Triangle^45^ thresholding in 2D, both of which also struggled with excessive merge errors. Methods labeled with asterisks were applied over 3D blocks of the volume, which in some cases resulted in the white “tiles” in some of the images.

### APEX2-Labeled Rodent Brain (ALRB)

The ALRB dataset (Figure 2 Panel F) is much larger than SRB, consisting of a 3948 × 1858 × 534 voxel volume imaged at a resolution of 1.2μm (1.9GB), which encompasses a coronal section of the rodent brain. To prepare this sample an 81 day old mouse was transcardially perfused with 4% paraformaldehyde, 2.5% glutaraldehyde in a 0.1 M sodium phosphate buffer solution. A 300μm coronal slice was prepared on a vibratome, reacted with DAB-Ni, then treated with 1% osmium tetroxide for 40 minutes, dehydrated through graded alcohols and acetonitrile as a transitional solvent, then embedded in Durcupan epoxy resin. The detailed protocol is described by Lam *et al*.^48^.

In addition to cells and vasculature, this dataset contains APEX2-labeled cells as a third class of structure to segment (Figure 2 Panels G & H). The APEX2-labeling was applied to a subpopulation of layer 4 cells. No ground-truth annotations exist for this volume, making a quantitative evaluation of the reconstructions impossible. We instead report properties of the segmented objects to compare the performance of FLoRIN reconstructions from 2D and 3D NDNT segmentations.

Both FLoRIN methods segmented 123,424 non-APEX2-labeled cells and 1,524 APEX2-labeled cells in ALRB, and the vasculature was segmented as a single connected component. Moreover, the reconstructions created by both FLoRIN methods were a Hausdorff distance of 1.614 × 10^−3^ apart. These findings indicate that 2D and 3D NDNT are able to discover approximately the same information, which FLoRIN then refines. As seen in Figure 2, the APEX2-labeled cells are visually distinct from the unlabeled cells and vasculature; the Filtering stage of the pipeline is able to distinguish between the two types. Such fine-grained distinctions are possible when the segmentation method is not dependent upon explicit training that may overlook a diversity of relevant features.

The FLoRIN and raw NDNT results are qualitatively better than other methods, as can be seen in Supplementary Figure S4. Where 2D and 3D NDNT were able to avoid artifacts in the, other methods with a similar result to NDNT were unable to overcome these *(e.g.*, Isodata^49^, Otsu^47^, and Li^50^ applied over 3D blocks, as well as Sauvola^44^). Other methods either could not produce useful segmentations or else severely under-segmented the volume.

### sSEM Volumes

Eliminating learning from the segmentation and reconstruction process potentially allows for generalization across different imaging modalities. Whereas the goal of segmenting and reconstructing μ CT X-ray volumes is to discover large structures and high-level information about the structure of the brain, sSEM images are of high enough resolution to follow individual processes (Figure 4 Panel A). The dataset introduced by Joesch *et al*.^51^ contains APEX2-positive processes with a sample preparation designed to enhance microstructures of interest for automatic sparse segmentation (Figure 4 Panel B). APEX2 labels starburst amacrine cells (SAC) in the dataset, thus we attempt to reconstruct the associated dendritic branches of these cells (Figure 4 Panels C-F).

**Figure 4.**
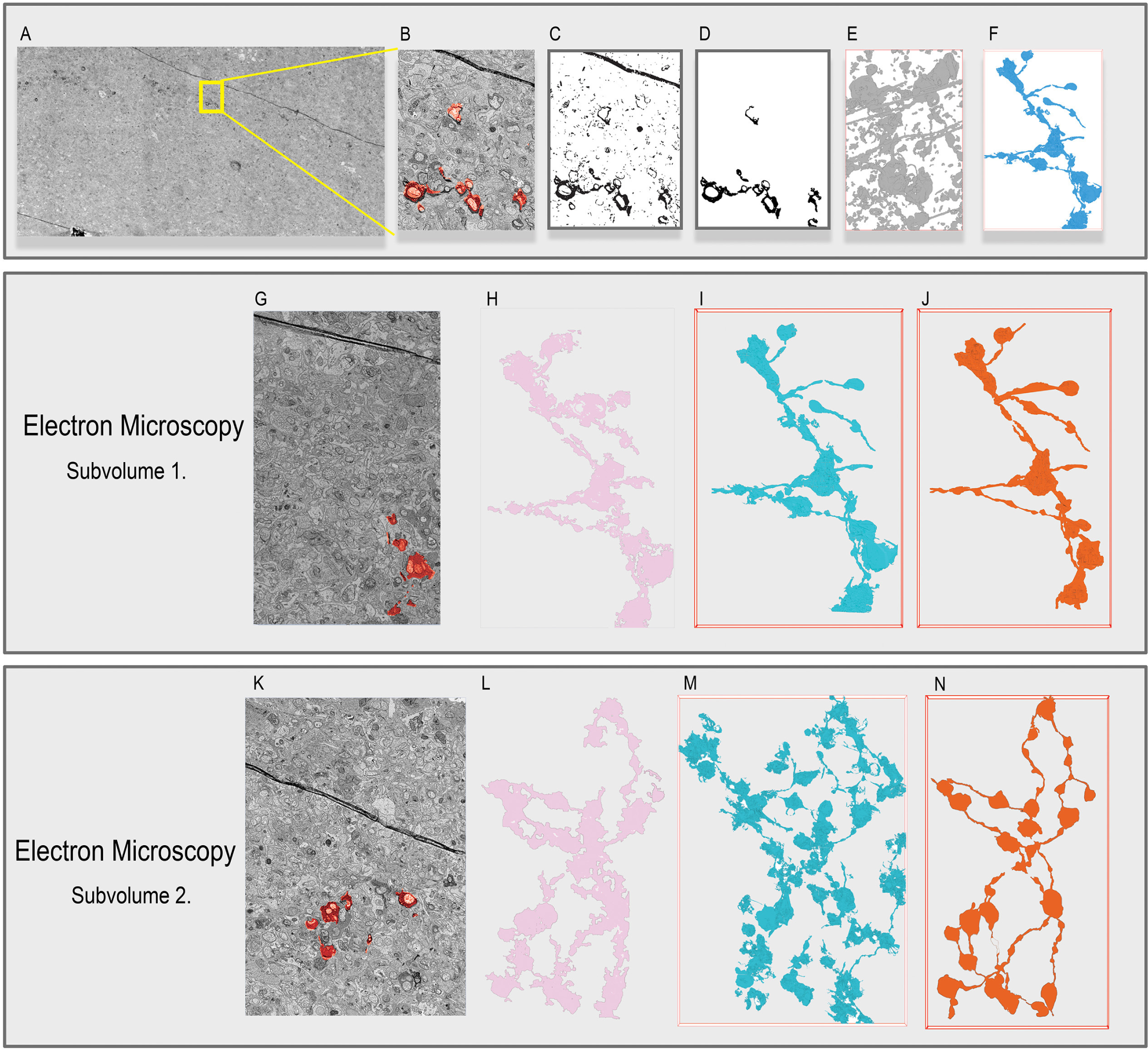
(A, B) The APEX2 positive sSEM sample contains a number of APEX-labeled SAC cells with annotated dendritic branches. The FLoRIN pipeline can conduct Segmentation and Filtering in 2D (C, D) or 3D (E, F). Segmentation (C, E) captures extraneous information, including non-APEX processes and artifacts, and these are removed in the Filtering stage (D, F). The middle panel shows (G) a sample of Subvolume 1 of the APEX2 positive sSEM sample, (H) the 2D reconstruction created by 2D FLoRIN, (I) the 2D reconstruction created by 3D FLoRIN, (J) ground-truth annotations of the APEX-labeled SAC cells. Both FLoRIN methods were able to reconstruct the annotated regions with many of the processes. While the 2D FLoRIN reconstruction incurred a number of merge errors on collapsing the reconstruction into a single plane, many of these issues are handled by 3D FLoRIN filtering connected components based on volumetric information. The bottom panel shows (K) a sample of Subvolume 2 of the APEX2 positive sSEM sample, (L) the 2D reconstruction created by 2D FLoRIN, (M) the 2D reconstruction created by 3D FLoRIN, (N) ground-truth annotations of the APEX-labeled SAC cells. Due to fewer artifacts in this subvolume, the two FLoRIN methods captured roughly the same information. Notably, though, 3D FLoRIN discovered additional pieces of the APEX2 positive process in this subvolume that were not present in the ground-truth data.

#### Subvolume 1

The first subvolume consisted of a 3,000 × 5,000 × 150 voxel region (4nm/pixel, 1.1GB) with substantial accompanying ground-truth annotations (Figure 4 Panel G). We processed this stack using FLoRIN in 2D and 3D, U-Net, 3D U-Net, and the learning-free methods listed in Supplementary Method: Non-Machine Learning Segmentation Method Evaluation. In every case, the methods attempted to segment and reconstruct APEX2-positive segments of SACs.

Figure 4 Panels H and I show the full 2D reconstructions created by FLoRIN from the 2D NDNT and 3D NDNT segmentations. A 2D reconstruction is the flattened version of the reconstruction that represents all of the segmented structures in a single plane. The 2D reconstruction is useful when studying SAC cells that are shallow along one dimension of the stack, allowing a unified view of the connections between cells, while a 3D reconstruction preserves depth information. Both reconstructions correspond well to the ground-truth, however the 3D NDNT reconstruction captured an additional APEX2-positive segment that was not included in the manual annotations.

As shown in Table 2, the FLoRIN reconstruction was the closest to the manual annotations, at a Hausdorff distance of 2.06 voxels. The raw 2D and 3D segmentations are also closer to the ground-truth than other thresholding results, with the remaining segmentations at least double and at most two orders of magnitude the distance. Additionally, FLoRIN was once again able to generate a reconstruction several orders of magnitude more quickly than U-Net and 3D U-Net could train on the sSEM data, and was roughly three times faster when making predictions.

**Table 2.** Performance comparison between FLoRIN, NDNT, U-Net models, and standard thresholding methods on Subvolume 1 and Subvolume 2 of the sSEM volume. U-Net results are the mean performance of 20 randomly-initialized nets. Asterisks indicate methods applied to small 3D blocks of the volume at a time. Hausdorff distances of ∞ indicate that the output did not include any positive pixels.

| Method | Subvolume 1 |  | Subvolume 2 |  |
| --- | --- | --- | --- | --- |
|  | Mean Hausdorff Distance | Runtime (Minutes) | Mean Hausdorff Distance | Runtime (Minutes) |
| **FLoRIN** | 2.06 | 29.64 | 1.54 | 73.29 |
| **NDNT** | 8.00 | 25.72 | 2.45 | 60.56 |
| **NDNT-2D** | 8.26 | 1.54 | 4.88 | 3.78 |
| **Isodata\***<sup>49</sup> | 126.79 | 2.36 | 87.99 | 5.30 |
| **Isodata**<sup>49</sup> | 122.70 | 0.24 | 82.97 | 0.58 |
| **Isodata 3D**<sup>49</sup> | 127.74 | 0.09 | ∞ | 0.26 |
| **Li\***<sup>50</sup> | 128.04 | 15.38 | 89.26 | 37.88 |
| **Li**<sup>50</sup> | 121.45 | 1.78 | 79.55 | 4.41 |
| **Li 3D**<sup>50</sup> | 123.31 | 0.09 | ∞ | 0.26 |
| **Local**<sup>54</sup> | 127.25 | 4.24 | 87.79 | 9.61 |
| **MGAC**<sup>46</sup> | 107.56 | 1044.51 | ∞ | 2555.33 |
| **Niblack**<sup>53</sup> | 125.09 | 3.39 | 87.83 | 7.37 |
| **Otsu\***<sup>47</sup> | 124.44 | 2.77 | 89.20 | 6.22 |
| **Otsu**<sup>47</sup> | 121.93 | 0.29 | 84.01 | 0.71 |
| **Otsu 3D**<sup>47</sup> | 124.02 | 0.08 | ∞ | 0.27 |
| **Sauvola**<sup>44</sup> | 103.96 | 3.37 | 57.48 | 7.60 |
| **Triangle\***<sup>45</sup> | 118.48 | 2.13 | 85.52 | 5.24 |
| **Triangle**<sup>45</sup> | 17.60 | 0.21 | 12.81 | 0.47 |
| **Triangle 3D**<sup>45</sup> | 124.60 | 0.09 | ∞ | 0.26 |
| **Yen\***<sup>52</sup> | 125.19 | 2.31 | 92.51 | 5.18 |
| **Yen**<sup>52</sup> | 54.65 | 0.20 | 8.60 | 0.48 |
| **Yen 3D**<sup>52</sup> | 127.79 | 0.09 | ∞ | 0.26 |
| **U-Net**<sup>33</sup> | 65.73 ± 10.81 | Train: 1413.73<br>Test: 80.47 | 24.56 ± 4.64 | Train: 1518.93<br>Test: 193.17 |
| **3D U-Net**<sup>34</sup> | 74.26 ± 12.47 | Train: 715.97<br>Test: 78.75 | 33.85 ± 2.15 | Train: 713.42<br>Test: 184.07 |

As can be seen in Figure 5, the FLoRIN and NDNT results capture the APEX2-positive region (the dark region in the raw image) completely and with good separation from surrounding noise. The Yen^52^ thresholding result was closest to the ground-truth after the FLoRIN and NDNT segmentation, but it could not establish an acceptable result for other modalities. The U-Net^33^ result was worse than the FLoRIN result, containing many merge errors and artifacts, and 3D U-Net^34^ was unable to segment the subvolume at all. Thresholding methods such as Isodata^49^, Li^50^, Otsu^47^, Niblack^53^, triangle^45^ and local^54^ thresholding, demonstrated poor thresholding and segmentation ability with output containing all microstructures, and not just the target APEX2 labeled process.

**Figure 5.**
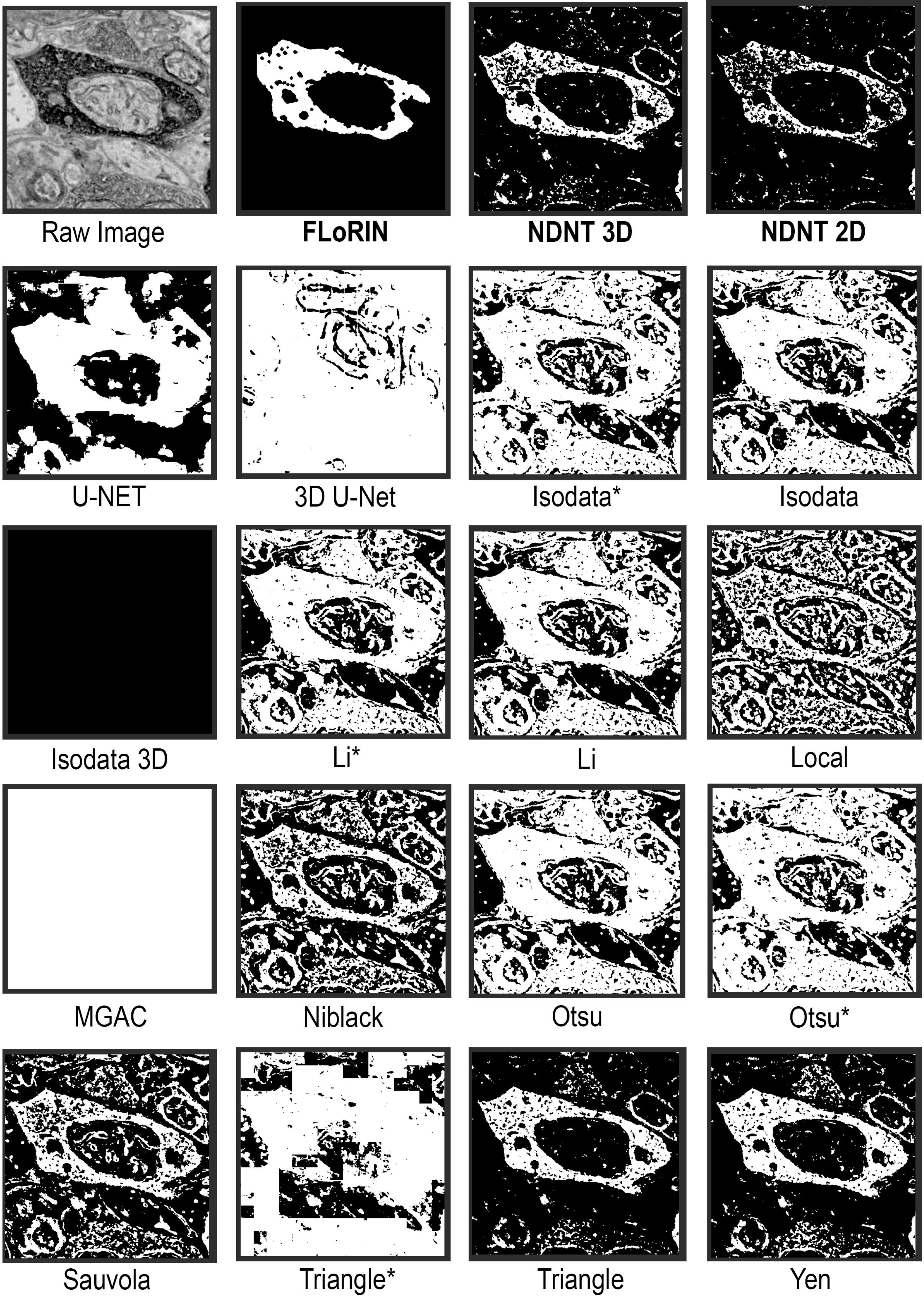
An example image from sSEM Subvolume 2 and the corresponding APEX2 labeled process segmentation results of various methods. The first row shows the result processed by the FLoRIN pipeline, and results of thresholding with NDNT 3D and 2D without any pre/post-processing. The 3D U-Net result was of poor quality, and the original U-Net result contains many merge errors. Similarly, other methods did not produce acceptable results. The Yen^52^ result comes close to being useful, but this method did not show the same performance in the other imaging modalities. Methods labeled with asterisks were applied over 3D blocks of the volume, which in extreme cases resulted in the white “tiles” in some of the images.

#### Subvolume 2

The second subvolume of the sSEM dataset consisted of 4,500 × 6,500 × 190 voxels (4nm/pixel, 4.4 GB), with large artifacts obscuring APEX2 positive processes, major grayscale shifts, and additional positive segments beyond the ground-truth annotations (Figure 4 Panel K). Once again, the 3D FLoRIN reconstruction was closer to the expert annotations than that of 2D FLoRIN (Figure 4 Panels L-M).

The 2D FLoRIN reconstruction had a final recall of 93.03%, however the reconstruction (Figure 4 Panel L) merges a number of the annotated features. The 3D FLoRIN reconstruction (Figure 4 Panel M), on the other hand, was able to separate microstructures and even discovered APEX2 positive processes overlooked by annotators (Figure 4 Panel N). As with Subvolume 1, 3D U-Net^34^ was unable to segment and reconstruct this dataset.

Similar to the Subvolume 1 results, FLoRIN and raw 3D and 2D NDNT segmentations were the closest to the ground-truth by at least a factor of two, as shown in Table 2. FLoRIN was also almost three times as fast as U-Net^33^and 3D U-Net^34^ generating a segmentation of the volume.

### Spectral Confocal Reflectance Microscopy Volumes

With μ CT X-ray and sSEM reconstructions, we demonstrated that FLoRIN is capable of reconstructing relevant structures in *ex-vivo* images. To determine if FLoRIN could also be applied to *in-vivo* data we also analyzed high-resolution optical images acquired from the live mouse brain. SCoRe is a recently developed technique that allows for precise label-free imaging of myelin in the live brain and in tissue samples^55^. SCoRe can be combined with fluorescence imaging in order to visualize patterns of myelination along stretches of single axons. In this modality, myelin patterns along single axons are captured by combining SCoRe (Figure 6 Panels C & D) with confocal fluorescence imaging in a transgenic mouse with YFP (Figure 6 Panels A & B) labeling on a subset of axons. FLoRIN was applied to this volume to reconstruct two independent myelinated axons from among a large number of visible intersecting background axons. 3D U-Net could not be applied to this problem because no annotations exist to train on SCoRe data.

**Figure 6.**
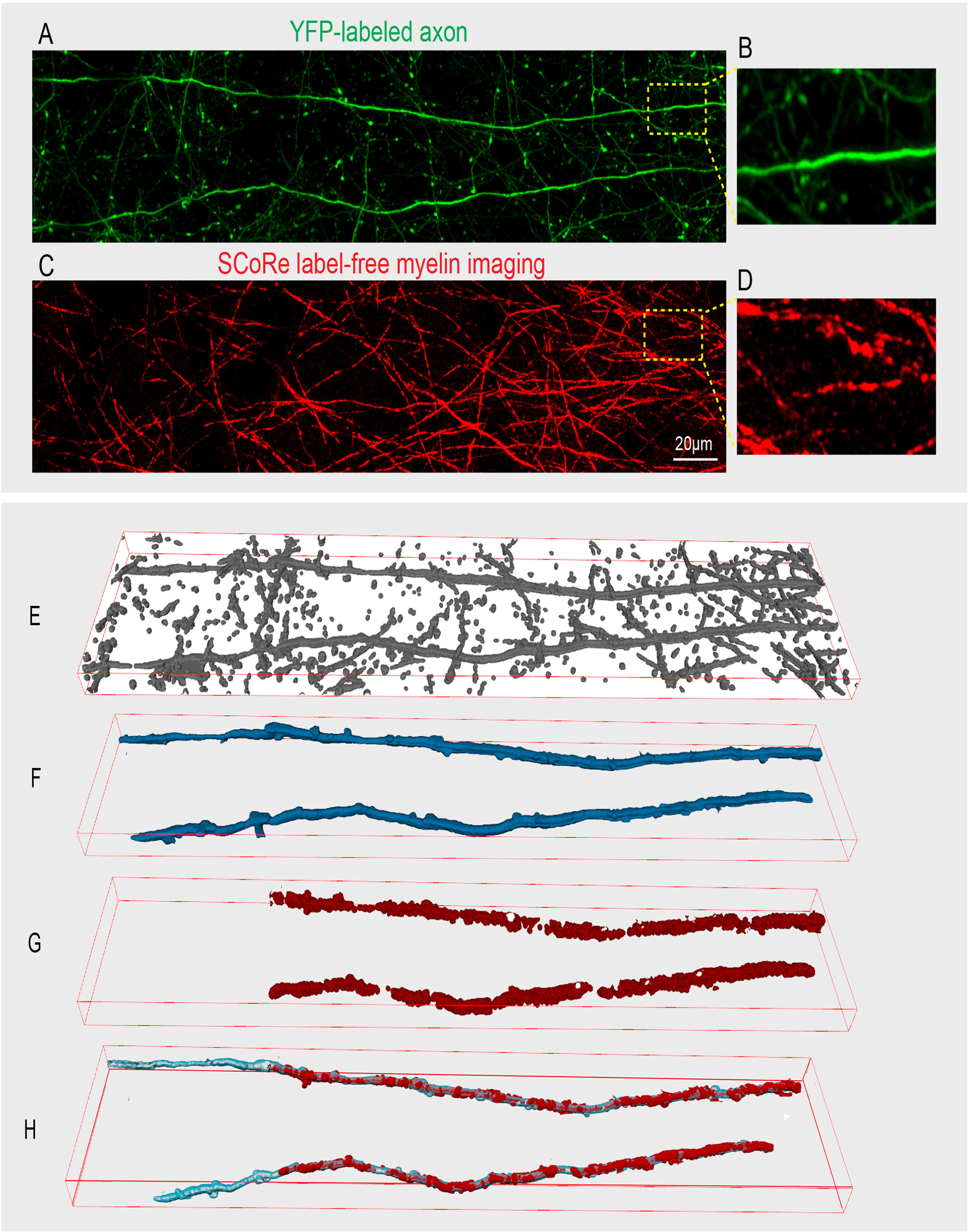
The SCoRE modality represents *in-vivo* (A-B) images captured from the cerebral cortex of a live mouse showing confocal fluorescence signals from Thy1-YFP labeled axons. (C-D) Label-free SCoRe images captured from the same region showing single myelinated fibers. (E) FLoRIN segmented axons generated using YFP signal. (F) Automated isolation and reconstruction of two YFP labeled axons within the volume. (G) corresponding SCoRe signals overlapping with the reconstructed axons. (H) Final reconstructed myelinated axons revealing unmyelinated and myelinated portions of the reconstructed axons.

We applied FLoRIN to a 311 × 66 × 21μ m (4.5 MB) SCoRe volume to reconstruct a single, contiguous myelinated axon. This volume contains a number of other axons that intersect with the axon of interest. Initially, the YFP channel was segmented by 3D NDNT (Figure 6 Panel E), and connected components were filtered to remove any unconnected segments (Figure 6 Panel F). Segments from the SCoRe channel were then acquired by only considering those overlapping the segmented portions of the YFP channel (Figure 6 Panel G). The final reconstruction (Figure 6 Panel H) combines the YFP and SCoRe channel segmentations.

FLoRIN using 3D NDNT revealed two long axons in the web of background axons, segmented as two connected components. Notably, FLoRIN was the only method able to follow both axons over a long distance in the YFP channel, the left part of the axon is not labeled with SCoRe, thus showing the specificity of FLoRIN for identifying overlap. These factors together indicate that FLoRIN reconstructions are able to discover far-reaching processes without training. FLoRIN created this reconstruction in 38.92s.

As can be seen in Supplementary Figure S5, 2D and 3D NDNT were the only methods to fully segment the intended axons with no merge errors. The only other methods to approach the performance of NDNT were Isodata^49^, Li^50^, and Otsu^47^ thresholding applied over 3D blocks at a time and Sauvola^44^ thresholding. These four methods also captured multiple brighter background axons that intersected with the intended axons, however, whereas NDNT was able to ignore these in both cases. All other methods severely over- or under-segmented the volume.

## Discussion

To understand the mechanisms of neural computation, access to high-quality segmentations and reconstructions of neuronal structures is paramount. FLoRIN demonstrably meets these needs and can do so in a fraction of the time needed to employ learning-based solutions. Whereas U-Net and other deep learning models require expert-annotated training data, which is time-consuming and costly to generate, FLoRIN is able to process new data without *a priori* information. This was demonstrated by the reconstructions of the ALRB and SCoRe volumes, which have no associated manual annotations. Moreover, FLoRIN bypassed the training time required to tune learning-based models by virtue of being learning-free, reducing both the computational time and the overall burden of model selection.

Across all datasets and modalities in this study, FLoRIN was able to create reconstructions faithful to expert-annotated ground-truth. In all cases, 3D NDNT-based FLoRIN produced results that were several factors closer to the ground-truth than U-Net trained on the ground-truth was able to produce. In the case of the SRB volume, the FLoRIN reconstruction was a factor of 30 closer to the ground-truth than the U-Net reconstructions. In the case of the sSEM volumes, however, 3D U-Net was unable to generate any meaningful results due to the similarity of APEX2-positive structures and other non-APEX-labeled features. FLoRIN reconstructed the APEX2-positive processes within both subvolumes and successfully filtered out false positives. Thus, FLoRIN outperformed a state-of-the-art deep learning model in multiple modalities.

Another key weakness of deep learning methods is the requirement that models be trained independently on examples of each microstructure of interest. FLoRIN, by contrast, is able to segment multiple classes of structures simultaneously: in each of the reconstructed μ CT X-ray volumes, the cells and vasculature were segmented in a single pass, and in the ALRB reconstruction, APEX-labeled cells were also captured. In this way it is able to handle segmentation errors, such as merge errors between different types of microstructure, by applying intermediate watershedding prior to labeling microstructures. While there are limitations to the simultaneous multi-class segmentations that arise when microstructures have high contrast with one another, this simply requires re-processing the image volume with a different threshold value or window size.

On top of outperforming U-Net, FLoRIN was applied to new volumes and new modalities without requiring extensive manual annotations for training. FLoRIN was successfully applied to segmenting a new μ CT X-ray volume, creating the first reconstruction of the ALRB volume without any prior information. Additionally, the FLoRIN statistical report provided initial data about the distribution of cells in the ALRB volume. This capability is not limited to previously-processed modalities: FLoRIN also created reconstructions of a SCoRe volume with no prior training, tracing far-reaching myelinated axons with no breaks. That FLoRIN was able to successfully reconstruct new volumes and modalities indicates that it is more generalizable than deep learning methods, which require additional training or complete re-training to effectively handle new datasets and modalities.

FLoRIN successfully solves the sparse segmentation and reconstruction problems by taking advantage of the inherent properties of sample preparations and by operating on local neighborhoods of voxels. The NDNT thresholding algorithm (Supplementary Algorithm S1) employed by FLoRIN is able to focus in on small regions, making it robust to grayscale shifts and distant noise that would otherwise confound global thresholding methods. Moreover, tissue preparation enhances the contrast of features of interest, in turn allowing for a stronger local threshold. NDNT can also account for volumetric information, allowing FLoRIN to process neural volumes as *volumes* rather than sequences of images. End-to-end processing in 3D can lead to greater biological fidelity, as demonstrated by the fact that 3D FLoRIN consistently created sparse reconstructions of higher quality than 2D FLoRIN.

Compared to existing thresholding methods, NDNT is robust to noise and generalizable across datasets and modalities. For each dataset in this study, we compared 2D and 3D NDNT segmentations and subsequent refinement during the Identification and Reconstruction stages of the FLoRIN pipeline against standard thresholding methods as described in Supplementary Method: Non-Machine Learning Segmentation Method Evaluation. In every case, NDNT segmentations were several factors to an order of magnitude closer to manual annotations than standard thresholding methods and active contours. This is due to the threshold value parameter: while other thresholding methods automatically select a threshold value based on global or neighborhood voxel values, NDNT allows the threshold value to be selected by a human through grid search, allowing an expert to refine the results based on domain knowledge. Tiling is also a contributing factor: processing small blocks or patches at a time reduced the impact of noise. Other methods that automatically select the threshold value are not necessarily improved using tiling, as shown in the quantitative results.

Expert-annotated ground-truth is costly, but necessary to make the best use of deep learning methods. In the absence of such data, we are forced to turn to learning-free methods for automated segmentation and reconstruction. As our reconstructions demonstrate, both NDNT-based FLoRIN outperforms deep learning in the data-starved regimes of μ CT X-ray and sSEM sparse segmentation by bypassing learning. Learning-free methods are generalizable across imaging modalities because they do not tune to specific features and noise, making them suitable for creating candidate reconstructions which may then be validated by experts. We suggest that FLoRIN be used for preliminary study and semi-automated annotation. Reconstructions created by FLoRIN can be used as a starting point for deciding how to study neural volumes: experts can gain a high-level view and choose regions to study at higher resolutions. Further study is possible by validating FLoRIN reconstructions — for example, examining each segmented microstructure alongside the original images and determining the class of structure it represents — and then training a deep learning model on the validated reconstructions. In this way, experts can make full use of the enormous amount of data being generated by new imaging technologies.

## Methods

### FLoRIN Framework

The FLoRIN pipeline is an open-source software package (https://github.com/CVRL/florin-scirep) for flexible, learning-free sparse segmentation and reconstruction of neural volumes. FLoRIN is divided into three stages, each of which use a series of learning-free image processing methods in either 2D or 3D: the Segmentation stage, Identification stage, and Reconstruction stage.

#### Stage 1. Segmentation

Starting with a set of raw images of a neural volume, the Segmentation stage optionally adjusts the grayscale histogram of the images, then thresholds the result. A number of standard grayscale adjustments are available, including histogram equalization, histogram re-scaling, and Wiener filters. Thresholding is carried out using the N-Dimensional Neighborhood Thresholding (NDNT) algorithm described in Supplementary Algorithm S1. It is possible to swap NDNT for any segmentation method during this stage, however we have found that NDNT consistently outperforms other learning-free methods. Operations in the segmentation stage are applied in 3D by default to take advantage of volumetric context, however FLoRIN may also fall back to 2D thresholding if needed (*e.g.*, for performance, to handle inconsistencies in 3D). The result of the Segmentation stage is a binarized version of the raw images that labels potential microstructures of interest.

#### Stage 2. Identification

The Identification stage takes the binary output of the Segmentation stage and optionally performs morphological operations before grouping connected components. Noise and artifacts may be addressed by removing small connected components, and combinations of dilations and erosions may be run to further clean the segmentation. The remaining connected components are then found and filtered into groups based on their geometric and grayscale properties. Connected components can be found in 3D to recover volumetric models directly, or else in 2D and stitched into 3D models. Supplementary Figure 1 provides an overview of the 2D filtering process. Information about the connected components is saved to a database for further review. Details of the resulting statistical report characterizing the anatomy of interest are described below.

#### Stage 3. Reconstruction

After Identification, FLoRIN saves each class of microstructure into an individual volume in the Reconstruction stage. Volumes can be output in a number of standard file types for compatibility with various postprocessing and rendering software. If a 2D reconstruction is desired, FLoRIN will collapse the computed volumes into a single plane and save the plane to file. Additionally, a statistical report is compiled from the database created during the Identification stage and can be modified to suit the content of the reconstruction.

#### Statistical Report

Along with the 3D reconstruction, FLoRIN outputs a Statistical Report that contains spatial, morphological, and grayscale information about each segmented microstructure. In addition to single-microstructure properties such as centroid, bounding box, volume, spatial orientation, and mean grayscale intensity, FLoRIN can compute statistics over sets of microstructures. Such statistical data allows for investigation into microstructure sizes, spatial distribution, or relationships between different types of microstructure. When evaluating the ALRB dataset, for example, we used the statistical report to compute the total number of discovered cells, average cell size, and average minimum intercellular distance. The statistical report allows for flexibility in how a neural volume is investigated. Often, the imaged volume is larger than the neural region of interest, as in the ALRB, sSEM, and SCoRe datasets evaluated in this study. In these cases, the data in the statistical report may be filtered to compute local statistics over the region of interest. Such a capability allows for multi-scale study of wide and targeted regions of the brain.

#### N-Dimensional Neighborhood Thresholding

Key to learning-free segmentation and reconstruction with FLoRIN is the NDNT algorithm, based on the method described by Bradley and Roth^41^. As shown in Supplementary Algorithm S1, NDNT computes the integral image over an arbitrary number of dimensions, allowing for thresholding to incorporate higher-dimensional contextual information such as depth in neural volumes. The threshold computation described by Bradley and Roth is then extended to operate across all dimensions of the computed integral image using the formula described by Tapia^42^.

The predecessor to FLoRIN, introduced in Joesch *et al.*^51^ used Otsu’s method^47^ to binarize neural images. Otsu thresholding uses a global value to determine the threshold, making it vulnerable to grayscale shifts and low signal-to-noise ratios. NDNT, by contrast, operates on a small neighborhood of voxels at a time, adaptively choosing a threshold for each voxel in the volume. As shown in Supplementary Figure S2, NDNT is tolerant to a variety of factors that confound global thresholding methods. Additionally, NDNT is significantly faster than global thresholding. We compared the running time against the previous version of the pipeline by segmenting and reconstructing the sSEM volume collected by Joesch *et al.*^51^ with FLoRIN and 3D

U-Net. As shown in Supplementary Table S1, FLoRIN is an order of magnitude faster than its predecessor due to the change in thresholding method and updates to the Identification stage. Based on both of these findings, we conclude that NDNT is primarily responsible for both the increased quality and speed of the FLoRIN pipeline.

### μ CT X-ray Data Collection

The tissue for the SRB dataset was acquired under animal procedures that were in accordance with NIH guidelines and protocols and approved by the institutional animal care and use committee at University of Chicago. The tissue for the ALRB dataset was acquired under animal procedures that were in accordance with NIH guidelines and protocols and approved by the institutional animal care and use committee at Allen Institute of Brain Science. Both of the SRB and ALRB image datasets were acquired at the Advanced Photon Source (APS) synchrotron beamline 32-ID-C at Argonne National Laboratory and Allen institute of Brain Science respectively. Both of the SRB and ALRB datasets were acquired at the Advanced Photon Source (APS) synchrotron beamline 32-ID-C at Argonne National Laboratory^56^ and Allen institute of Brain Science respectively. To minimize the number of artifacts in the tomography the beamline used no optics to filter the X-ray beam. Data acquisition was performed in propagation-based phase contrast mode with 350mm between the detector and sample with a pink beam (δE/E = 10-2) at 25keV energy.

Each tomogram was acquired with 1601 projections at different rotation angles equally spaced from 0 to 180 degrees. The projections were acquired with an X-ray to visible light microscope that consists of a scintillator followed by a light microscope. The light is then focused by a Mitutoyo High Resolution 5x objective lens on a 1920 × 1200 pixel array detector (Point Grey, GS3-U3-23S6M-C). The numerical aperture of the lens was .21 giving a resolving power of 1.31 μm, the pixel size of the camera was 1.17 μm and the resulting field of view was 2.25 × 1.41mm^2^. Each tomogram took three minutes to be acquired and two minutes to be reconstructed on the Cooley HPC cluster at Argonne National Laboratory. The automatization of the process was done by AuTomo^56^ and the final reconstruction by the gridrec algorithm of TomoPy^57^. No phase retrieval was performed.

### sSEM Data Collection

Datasets were acquired as described by Joesch *et al.*^51^. Animals were used in accordance with NIH guidelines and protocols approved by Institutional Animal Use and Care Committee at Harvard University. In short, retinas of ChaT-cre mice were anesthetized and infected with rAAV2 expressing APEX2NES under the CAG promoter. 4 weeks following infection, the retina was dissected and fixed with PFA and glutaraldehyde. Following aldehyde fixation, the tissue was washed and reacted with DAB to reveal sites of peroxidase activity. The DAB polymer was subsequently reduced with sodium hydrosulfite, stained with osmium, dehydrated and infiltrated with Durcupan resin. The cured blocks were trimmed and serially sectioned (~30 nm) using a custom tape collection device (ATUM^58^) attached to a commercial ultramicrotome. Sections were collected on plasma-treated carbon-coated polyamide (Kapton, Sheldahl) 8-mm-wide tape and post-stained with uranyl acetate in maleate and with lead citrate. An automated protocol to locate and image sections on the wafers was used^58^ with a Sigma scanning electron microscope (Carl Zeiss), equipped with the ATLAS software (Fibics). Images were acquired using secondary electron detection. Data was then aligned using non-affine alignment through the FijiBento alignment package (https://github.com/Rhoana/FijiBento^51^).

### SCoRe Data Collection

All animal procedures were in accordance with NIH guidelines and protocols and approved by the institutional animal care and use committee at Yale University. Postnatal day 60 male transgenic mice (Thy1-YFPh, Jax# 033782) were used for *in-vivo* SCoRe and fluorescence imaging. Mice were anesthetized with Ketamine/Xylazine and a 3mm cranial window was prepared over the somatosensory cortex as previously described^55^. The mice were immediately imaged on a confocal microscope (Leica SP5) with a 20x water immersion objective (1.0NA Leica). For SCoRe imaging, the confocal reflected signals from 488nm, 561nm, and 633nm wavelength lasers were combined into a single image in order to visualize the myelin sheath in a label free fashion^55^. The fluorescence signal from Thy1-YFP labeled axons was collected sequentially in the same cortical region using 488nm wavelength excitation.

### Evaluation Metrics

The reconstructions used in this study were evaluated against expert annotations using Hausdorff distance, a similarity measure that determines the maximum distance between any point in the reconstruction and the nearest point in the annotations. Details about this metric may be found in Supplementary Methods: Evaluation Metrics.

## Data Availability

The segmentations and reconstructions generated during the current study are available from the corresponding author on reasonable request. The SRB dataset is available for download from the XBrain project (https://github.com/neurodata/xbrain). The sSEM volume along with manual annotations, has been published to Dryad^59^. The SCoRe volume used in this study is available from Jaime Grutzendler on reasonable request.

## Acknowledgements

Ali Shahbazi, Jeffery Kinnison, Walter Scheirer, Rafael Vescovi, and Narayanan Kasthuri were supported by the Office of the Director of National Intelligence (ODNI), Intelligence Advanced Research Projects Activity (IARPA) under contract #D16PC00002. Marc Takeno, and Nuno M. da Costa were supported by IARPA via Department of Interior/ Interior Business Center (DoI/IBC) contract number #D16PC00004. The views and conclusions contained herein are those of the organizers and should not be interpreted as necessarily representing the official policies, either expressed or implied, of ODNI, IARPA, DoI/IBC, or the U.S. Government. The U.S. Government is authorized to reproduce and distribute reprints for governmental purposes notwithstanding any copyright annotation therein. Equipment was generously donated by the NVIDIA Corporation, and made available by the National Science Foundation (NSF) through grant #CNS-1629914. This research used resources of the Argonne Leadership Computing Facility, which is a DOE Office of Science User Facility supported under Contract DE-AC02–06CH11357.

## Author Contributions Statement

Ali Shahbazi and Jeffery Kinnison developed the FLoRIN framework, carried out all segmentation and reconstruction processes, wrote the main manuscript text, and prepared Figure. 1–6. Walter Scheirer designed and led the research. Rafael Vescovi, Ming Du, and Narayanan Kasthuri provided the SRB μ CT X-ray data used in this study and contributed the description of this dataset. Marc Takeno, Hongkui Zeng, and Nuno da Costa provided the ALRB μ CT X-ray data used in this study and contributed the description of this dataset. Maximilian Joesch provided the sSEM data used in this study and contributed the description of this dataset. Robert Hill and Jaime Grutzendler provided the SCoRe data used in this study and contributed the description of this dataset. All authors reviewed the text of the manuscript.

## Additional information

The authors declare no competing interests.

